# Inferring multi-scale neural mechanisms with brain network modelling

**DOI:** 10.1101/157263

**Authors:** Michael Schirner, Anthony Randal McIntosh, Viktor K. Jirsa, Gustavo Deco, Petra Ritter

## Abstract

The neurophysiological processes underlying non-invasive brain activity measurements are not well understood. Here, we developed a novel connectome-based brain network model that integrates individual structural and functional data with neural population dynamics to support multi-scale neurophysiological inference. Simulated populations were linked by structural connectivity and, as a novelty, driven by electroencephalography (EEG) source activity. Simulations not only predicted subjects’ individual resting-state functional magnetic resonance imaging (fMRI) time series and spatial network topologies over 20 minutes of activity, but more importantly, they also revealed precise neurophysiological mechanisms that underlie and link six empirical observations from different scales and modalities: (1) slow resting-state fMRI oscillations, (2) spatial topologies of functional connectivity networks, (3) excitation-inhibition balance, (4, 5) pulsed inhibition on short and long time scales, and (6) fMRI power-law scaling. These findings underscore the potential of this new modelling framework for general inference and integration of neurophysiological knowledge to complement empirical studies.

## Introduction

Neural activity is pervaded by dynamic patterns on multiple spatiotemporal scales, whose mutual relationships and significance for function and behaviour are unclear. Empirical approaches to characterizing the mechanisms that govern brain dynamics often rely on the simultaneous use of different acquisition modalities. These data can be merged using statistical models, but the inferences are constrained by information contained in the different signals, rendering a mechanistic understanding of neurophysiological processes elusive. Brain simulation is a complementary technique that enables inference on model parameters that reflect mechanisms that underlie emergent behaviour, but that are hidden from direct observation.

We developed a novel type of brain network model, dubbed ‘hybrid model’, where each subject’s source-modelled EEG data was used to drive local dynamics in a large-scale connectome-based model. Resulting hybrid models reproduced ongoing subject-specific fMRI time series over a period of 20 minutes and a variety of other empirical phenomena (Figure 1). In contrast to previous brain network models that used noise as input, hybrid models are driven by EEG source activity and therefore simultaneously incorporate structural and functional information from individual subjects (Figure 2). The injected EEG source activity serves as approximation of excitatory synaptic input currents (EPSCs), which helped increase the biological plausibility of generated model activity (Atallah & Scanziani, 2009; Buzsáki et al., 2012; Nunez & Srinivasan, 2006). Individualized hybrid models yielded predictions of ongoing empirical subject-specific resting-state fMRI time series (Figures 3). Additionally, spatial topologies of fMRI functional connectivity networks (Figure 4), E/I balance of synaptic input currents, α-rhythm mediated pulsed inhibition of population activity on short (Figure 5), and on long time scales (Figure 6), and fMRI power-law scaling were reproduced (Figure 7). More importantly, our subsequent analysis of intrinsic model activity revealed neurophysiological processes that could explain how brain networks produce the aforementioned signal patterns (Figures 5 to 7). That is, simulation results not only predicted ongoing subject-specific resting-state fMRI time series and several empirical phenomena observed with invasive electrophysiology methods, but more importantly, they also show how the network interaction of neural populations lead to the emergence of these phenomena and how they are connected across multiple temporal scales in a time scale hierarchy.

**Figure 1.**
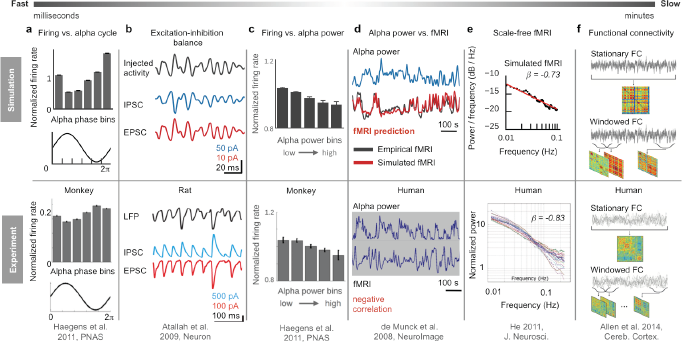
Overview of six empirical phenomena on different temporal scales reproduced by the hybrid model. In addition to reproducing empirical phenomena, model activity also reveals neurophysiological mechanisms underlying their emergence (Figures 5 to 7). (a) Neuron firing is related to the phase of alpha waves (8-12 Hz) (Haegens et al., 2011). During peaks of alpha waves neurons fire the least, while they fire with maximum rate during troughs. Behavioural studies indicate that this effect is related to neural information processing, which is formulated in ‘gating by inhibition’ and ‘pulsed inhibition’ theories (Jensen & Mazaheri, 2010; Klimesch et al., 2007). (b) Our simulation results indicate that this relationship between firing and alpha phase is related to the well-established observation of ongoing balancing of neural excitation and inhibition (Atallah & Scanziani, 2009; Isaacson & Scanziani, 2011; Okun & Lampl, 2008). The empirically observed relationship between local field potentials (LFPs), EPSCs and IPSCs is reproduced in hybrid models as a result of source activity injection and inhibitory population activity. Model activity indicates that the inhibitory effect of alpha (a) is related to ongoing E/I-inhibition balance mediated by inhibitory populations (Figure 5). (c) On a longer time scale (<0.25 Hz) neuron firing rates and task performance are inversely related to alpha power (Haegens et al., 2011), which was also reproduced by the model. (d) More importantly, simulation results indicate that the inverse relationship between firing and alpha power is the underlying cause of the inverse relationship between fMRI and alpha power oscillations, which is a well-established finding from simultaneous measurements of electric and hemodynamic neural activity (de Munck et al., 2008; Feige et al., 2005; Goldman et al., 2002; Moosmann et al., 2003). In particular, the model shows a direct mechanism that transforms alpha power oscillations into fMRI oscillations, which constitutes the first concrete hypothesis on the relationship between electric and hemodynamic resting-state oscillations (Figure 6). Importantly, the model was able to predict ongoing subject-specific resting-state fMRI time series and corresponding spatial network patterns of 15 subjects on the basis of their individual EEG and structural brain connectivity (Figure 3). (e) Relatedly, the model also explains the emergence of scale-free fMRI power spectra (He, 2011) as a result of global network interaction (Figure 7) and predicts f) individual functional connectivity matrices (Allen et al., 2014) computed over long and short time windows (Figure 4). The ability of the hybrid model to infer precise neurophysiological mechanisms that give rise to empirical phenomena and to link the involved mechanisms and signal patterns across different scales and neuroimaging modalities makes it a potentially valuable tool for neuroscience research.

**Figure 2.**
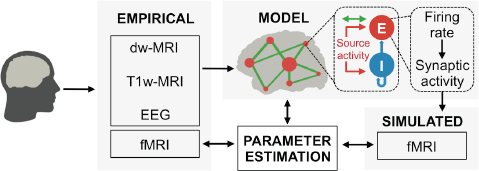
Hybrid modelling framework. Hybrid brain network models are constructed from diffusion-weighted MRI tractography results and region-parcellations obtained from T1-weighted MRI. In contrast to conventional models, hybrid models are injected with EEG source activity time series of the same subject, instead of noise. By tuning model parameters, predicted fMRI time series are fit to each subject’s empirical fMRI time series, which were simultaneously acquired with EEG. Injection of EEG source activity enabled better prediction of subject-specific fMRI time series compared to noise-driven brain network models (Figure 3). 5-fold cross-validation was performed to guard against overfitting. At each node (small red circles) of the large-scale network (green lines) are local networks of excitatory (E) and inhibitory (I) neural cell population models that are driven by EEG source activity (red arrows). Population models represent the activity of individual brain regions and are globally coupled by structural connectomes (green arrows) that represent the heterogeneous white matter coupling strengths between different brain areas. Firing rates and synaptic activity time series underlying fMRI predictions are analysed to identify how neural population activity and network interaction relates to observable neuroimaging signals. See also supplementary movie 1 for a visualization of brain network model construction and exemplary results from hybrid model simulations.

**Figure 3.**
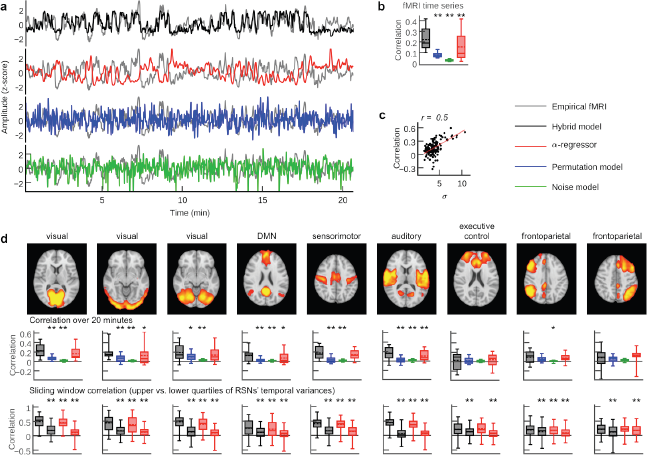
Person-specific fMRI time series prediction. (**a**) Example time series of the hybrid model and the three control scenarios. (**b**) Box plots of average correlation coefficients between all simulated and empirical region time series (20.7 min) for each subject (n = 15; values for the α-regressor were inverted for illustration purposes). (**c**) Scatter plot of RSN time course standard deviation (s.d.) versus prediction quality. Dots indicate the s.d. of each of the nine RSN time courses in each of the 15 subjects versus the prediction quality of fMRl time series for all regions underlying the respective RSN. Predictions are better when the corresponding RSN time course has a high s.d., i.e., when the RSN contributes a higher variance to the overall JMRl signal. (**d**) Box plots compare prediction quality during upper versus lower quartile of epoch-wise RSN time course s.d.s. Upper row: spatial activation patterns of nine RSNs. Middle row: correlation coefficients between RSN temporal modes and hybrid model simulation results and the three control setups. Lower row: sliding window (length = 100 fMRl scans = 194 s; one fMRl scan step width) correlations for the upper (first and third boxplot per panel) and lower quartiles (second and fourth boxplot per panel) of window-wise RSN temporal mode for the hybrid model and the α-regressor (n = 2010 epochs). Asterisks indicate significantly increased prediction quality of the hybrid model compared to control conditions in one-tailed Wilcoxon rank sum test (*p <0.05, **p <0.01). Additionally, all hybrid model correlations in (**b**) and (**d**) were tested for the null hypothesis that they come from a distribution whose median is zero at the 5 % significance level. All tests rejected the null hypothesis of zero medians except for RSN correlations over 20 minutes for the executive control and the frontoparietal networks (middle row).

**Figure 4.**
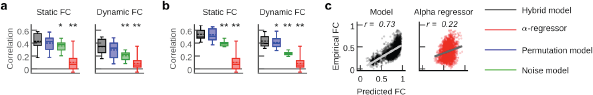
Functional connectivity prediction. In contrast to the α-regressor, the hybrid model concurrently predicts fMRI time series (Figure 3) and the spatial topology of fMRI networks. (**a, b**) Box plots show correlation coefficients between predicted and empirical FC for the three model types and the α-regressor (n = 15 subjects). FC was computed for long epochs (static FC; computed over 20.7 min) and short epochs (dynamic FC; average sliding window correlation; 100 fMRI scans window length; one fMRI scan step width). The hybrid model yields significantly higher FC correlations than noise model and α-regressor. Results were compared for the parameter set that generated the best fMRI time series prediction (**a**) and the parameter set that yielded the best FC predictions for each subject (**b**). (**c**) Scatter plots compare empirical and simulated average region-wise FC for hybrid model simulations and the α-regressor. Dots show all average region-wise static FC values over all subjects. Asterisks indicate significantly increased prediction quality of the hybrid model compared to control conditions in one-tailed Wilcoxon rank sum test (*p <0.05, **p <0.01).

**Figure 5.**
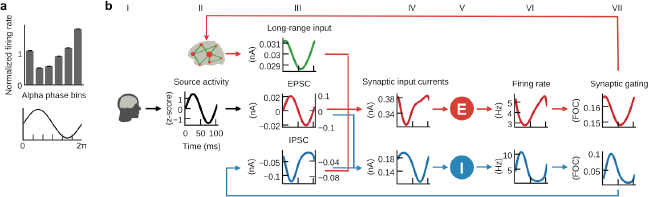
E/I balance generates pulsed inhibition. Hybrid models reproduced the invasively observed rhythmic inhibition of firing relative to α-cycle phase (Haegens et al., 2011). (**a**) Histogram of population firing rates divided into six bins according to α-cycle segments and normalized relative to the mean firing rate of each cycle. Population firing rates were highest during the trough and lowest during the peak of α-cycles. (**b**) Grand average waveforms of population inputs and outputs time locked to α-cycles of injected EEG source activity (black, column II). Shaded areas indicate standard errors of the mean; left and right axes denote input currents to excitatory and inhibitory populations, respectively. Inhibitory population inputs (blue, column IV) were dominated by EPSCs (red, column III). Consequently, firing rates and synaptic gating of inhibitory populations (blue, columns VI and VII) closely followed source activity shape. As inhibitory populations revert the effect of their input currents, the shape of resulting inhibitory postsynaptic currents (IPSCs; blue, column III) was inverted to source activity shape. Further reproducing empirical observations (Atallah & Scanziani, 2009; Xue et al., 2014), the amplitude of IPSCs at excitatory populations was several times larger than EPSCs. Accordingly, excitatory population input was inverted relative to the α-cycle, leading to rhythmic inhibition of firing rates (red, column VI) near the peak of the alpha cycle.

**Figure 6.**
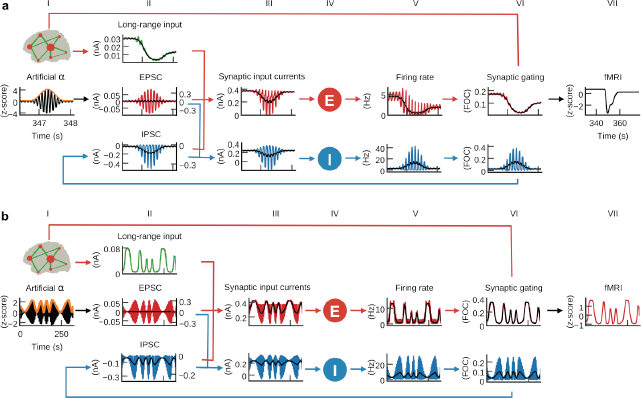
α-power fluctuations generate fMRI oscillations. Grand average waveforms of population inputs and outputs. (**a**) Hybrid models were injected with artificial α-activity consisting of 10 Hz sine oscillations that contained a single brief high power burst (black, column I; orange: signal envelope). While positive deflections of the α-wave generated positive deflections of inhibitory population firing rates, large negative deflections were bounded by the physiological constraint of 0 Hz (blue, fifth column; black: moving average). That is, inhibitory populations rectified input α-oscillations such that only the positive half¬wave had an inhibitory downstream effect (blue, column VII and II). As a result, average per-cycle feedback inhibition increased for increasing α-power, and consequently the average firing rates, synaptic gating and ultimately fMRI signals (red, column V, VI and VII) decreased. (**b**) Hybrid models were injected models with 10 Hz sine waves where ongoing power was modulated similar to empirical α-rhythms (0.01-0.03 Hz). Similarly to (**a**), but for a longer time frame, inhibitory populations rectified negative deflections, which introduced the α-power modulation as a new frequency component into firing rates and fMRI time series.

**Figure 7.**
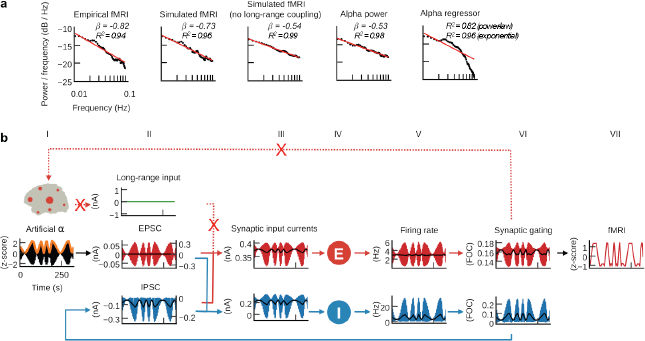
Long-range coupling controls fMRI power-law scaling. (**a**) Power spectral densities (straight-line fits are for illustration purposes only; scale-invariance was determined using rigorous model selection criteria, see Methods). Previous results indicated that the model transformed ongoing EEG source power fluctuation into fMRI oscillations. However, α-power time series had a considerably flatter power spectrum slope (β_α-band_ =-0.53, β_wide-band_ =-0.47) than fMRI. Simulation results that were obtained for deactivated large-scale coupling had a similarly flat power spectrum (β_zeroG_ =-0.54). Parameter space exploration suggests that the emergence of scale-free fMRI power spectra depends on the proper balancing of recurrent large-scale excitation with local inhibition (Figure 7—figure supplement 1 and Figure 7—figure supplement 2). When large-scale coupling was absent or inadequately balanced, prediction quality decreased and models produced flatter power spectra or no scale invariance at all. (**b**) As in Figure 6b, but with disabled large-scale coupling. In contrast to Figure 6b, the resulting firing rates, synaptic gating and fMRI waveforms showed no frequency-amplitude dependence. Comparison of model dynamics between both scenarios suggests that gradually accumulating self-reinforcing excitation through white-matter coupling leads to frequency-dependent amplification that augments slower oscillations more than faster oscillations, which results in the emergence of scale-free fMRI power spectra.

Resting-state fMRI studies identified so-called “resting-state networks” (RSNs), which are widespread networks of coherent activity that spontaneously emerge across a variety of species in the absence of an explicit task (Biswal et al., 1995; Fox & Raichle, 2007; Raichle et al., 2001). Despite correlations between fMRI and intracortical recordings (He et al., 2008; Logothetis et al., 2001), EEG (Becker et al., 2011; Goldman et al., 2002; Mantini et al., 2007; Moosmann et al., 2003; Ritter et al., 2009) and magnetoencephalography (Brookes et al., 2011; de Pasquale et al., 2010) the link between RSNs and electrical neural activity is not fully understood. Electrical neural activity is dominated by oscillations in the α-band, which is rhythmic activity in the 8 to 10 Hz frequency range first discovered by Hans Berger in 1929 (Berger, 1929). A growing body of research suggests that information processing, attention, perceptual awareness, and cognitive performance are rhythmically modulated by α-power and phase (Busch et al., 2009; Klimesch, 1999; Mathewson et al., 2009). The roles of α-rhythms for mediating top-down control, timing of oscillations and directing attention by blocking task-irrelevant pathways are central to prevailing hypotheses termed ‘gating by inhibition’ and ‘pulsed inhibition’ (Jensen & Mazaheri, 2010; Klimesch et al., 2007). Interestingly, intracellular recordings showed that inhibitory events are inseparable from excitatory events, resulting in an ongoing excitation-inhibition balance (E/I balance) (Isaacson & Scanziani, 2011; Okun & Lampl, 2008). The significance of the α-rhythm is underscored by strong negative correlations between its ongoing power fluctuation and resting-state fMRI amplitude fluctuation (de Munck et al., 2008; Feige et al., 2005; Goldman et al., 2002; Moosmann et al., 2003). Lastly, despite wide-spread interest in critical dynamics (Bak, 2013) the emergence of power-law scaling, a signal pattern that is ubiquitous in nature and commonly observed in neural activity, is unclear (Beggs & Timme, 2012; Marković & Gros, 2014).

To illustrate the potential of this framework for inference of neurophysiological processes we show inferred mechanisms for three different empirical phenomena and how they relate to other well-established neural signal patterns (Figure 1). Upon finding that the hybrid model predicts fMRI activity, we first sought to identify how injected EEG drove the prediction of subject-specific fMRI time series, which led us to a mechanism that transformed α-power fluctuations of injected EEG source activity into fMRI oscillations, which may explain the empirically observed correlation between EEG and fMRI. Consequently, we asked how the inhibitory effect of α-power oscillations was created, which led to the identification of an inhibitory effect on the considerably faster timescale of α-phase fluctuations. As these inhibitory effect of alpha rhythms on short and long time scales were mediated by the interaction of local populations, but prediction quality decreased when large-scale coupling was deactivated, we interrogated the model for the influence of structural coupling on the emergence of fMRI oscillations and found that global coupling amplified brain oscillations in a frequency-dependent manner, which facilitated the emergence of power-law scaling. Starting with fast-scale effects, our first model outcome accounts for the invasively observed inverse relationship between neuronal firing and α-rhythm phase by identifying a mechanism that explains this ‘pulsed inhibition’ with ongoing E/I balance. The second model outcome posits a neural origin of fMRI RSN oscillations by identifying an explicit mechanism that transforms ongoing α-power fluctuations into slow fMRI oscillations, which also explains the empirically observed anti-correlations between α-power and fMRI time series. Our third model outcome indicates that scale invariance of fMRI power spectra results from self-reinforcing feedback excitation via large-scale white-matter coupling, which leads to frequency-dependent amplification of neural oscillations.

## Results

### Hybrid models predict subject-specific fMRI time series

The brain network models used here are dynamical systems where individual brain areas are simulated by neural mass models that are coupled by weighted structural coupling strength estimates obtained from diffusion-weighted MRI white-matter tractography. Neural mass models are derived from networks of spiking neuron models to capture their essential modes of behaviour based on mean-field approximation techniques (Deco et al., 2008). In contrast to previous brain network models that used noise as input’ the neural mass models of our ‘hybrid’ model are driven by EEG source activity that was simultaneously acquired with fMRI (Figure 2). Simulation results predicted a considerable part of the variance of ongoing resting-state fMRI time series (Figure 3) and spatial network topologies (Figure 4). Furthermore, models that were fitted to subject-specific fMRI time series reproduced a variety of empirical phenomena observed with EEG and invasive electrophysiology (Figure 1) and, more importantly, simulation results revealed mechanistic explanations for the emergence of these phenomena (Figures 5, 6 and 7)

We constructed individual hybrid brain network models for 15 human adult subjects using each subject’s own structural connectomes and injected each with their own region-wise EEG source activity time courses. Using exhaustive searches, we tuned three global parameters for each of the 15 individual hybrid brain network models to produce the highest fit between each of the subject’s empirical region-average fMRI time series and corresponding simulated time series. The first parameter scales the strength of structural coupling and the second and third parameters scale the strengths of EEG source activity inputs injected into excitatory and inhibitory populations, respectively. To compare the quality of fMRI predictions, the hybrid model simulation results were compared with three control scenarios: *(i)* a brain network model where local dynamics were driven by noise, *(ii)* a variant of the hybrid model that used random permutations of the region-wise EEG source activity time series and *(iii)* a statistical model where ongoing α-band power fluctuation of EEG source activity was convoluted with the canonical hemodynamic response function (henceforth called α-regressor). The first two controls are brain network models and the third is inspired by traditional analyses of empirical EEG-fMRI data.

Visual inspection of example time series showed good reproduction of characteristic slow (<0.1 Hz) RSN oscillations by the hybrid model and the α-regressor (albeit inverted for the latter), but poor reproduction of temporal dynamics in the case of noise and random permutations models (Figure 3). To quantify prediction quality, we compared for each subject the average correlation coefficients over all 68 regions of the brain parcellation between all simulated and empirical fMRI time series for each of the four scenarios (i.e., hybrid model and the three control setups). Predictions from the hybrid model correlated significantly better with empirical fMRI time series than predictions from the two random models and the α-regressor (Figure 3b). For the hybrid model, five-fold cross-validation showed no significant difference of prediction quality between training and validation data sets (two-tailed Wilcoxon rank sum test, *p* = 0.71, *t* = 0.54, Cohen’s *d* = 0.013) and between validation data sets and prediction quality for the full time series (two-tailed Wilcoxon rank sum test, *p* = 0.42, *t* =-0.2, Cohen’s *d* =-0.036).

To estimate the ability of the four scenarios to predict the time courses of different commonly observed RSNs we performed a group-level spatial independent component analysis (ICA) of the empirical fMRI data. Next, we computed average correlation coefficients between each subject-specific RSN time course and the model regions at the position of the respective RSN. As in the case of region-wise fMRI (Figure 3b), correlation coefficients of the hybrid model were significantly larger than the control network models for most RSNs (Figure 3d). The sliding-window analyses showed that prediction quality varied over time, regions and subjects: window-wise prediction quality was highly correlated with the standard deviation of RSN temporal modes (Figure 3c, d). That is, the higher the variance contributed to overall fMRI activity by an RSN in a given subject and time window, the better the prediction of empirical fMRI. As a consequence, epochs in the upper quartile of RSN s.d.s were significantly better predicted than epochs in the lower quartile (Figure 3d). In order to assess subject-specificity of fMRI time series predictions we correlated all simulation results of each subject (i.e. for every tested parameter combination) also with the empirical fMRI activity of all other subjects. We found that the maximum correlation coefficients over all tested parameters were significantly larger when empirical and simulated data sets belonged to the same subject compared to when they came from different subjects (p <10-4, Wilcoxon rank sum test).

Next, we estimated the ability of all four setups to predict the spatial topology of empirical fMRI networks. In contrast to time series prediction, the α-regressor showed low correlations with empirical functional connectivity (FC). Compared to the α-regressor, all three model-based approaches provided significantly better predictions of subjects, individual long-term FC and short-term FC (Figure 4). Furthermore, hybrid model simulation results correlated significantly better with empirical network topology than predictions obtained from the conventional noise-driven model (Figure 4a,b). Interestingly, correlations for hybrid and random permutation models were effectively the same, likely because the large-scale network dynamics, enabled by structural coupling, that drive the emergence of FC would be relatively preserved when permuting injected activity. Prediction of group-average FC (all pairwise FC values averaged over all subjects) was better for the hybrid model compared to the α-regressor (Figure 4c).

### E/I balance generates pulsed inhibition

After fitting the individual hybrid models for each of the 15 subjects, we analysed the local population activity to infer neurodynamic mechanisms underlying predicted fMRI time series. Our first observation was that on the fast time scale of individual α-cycles (~100 ms) the optimized hybrid model reproduced the inverse relationship between α-phase and firing rates observed in invasive recordings (Haegens et al., 2011) (Figure 5a). To investigate these fast-acting dynamics related to α-phase, we computed grand average waveforms of modelled synaptic inputs, population firing rates, and synaptic gating time-locked to the zero-crossings of α-cycles. Resulting waveforms illustrate the relation between α-oscillations and neural firing and how the ongoing balancing of rhythmic excitatory and inhibitory inputs generated pulsed inhibition (Figure 5b).

As a result of optimizing the three model parameters, injected EPSCs dominated the inputs to inhibitory populations and the sum of synaptic input currents to inhibitory populations closely followed the shape of EPSCs. Also, because of the monotonic relationship between input currents and output firing rates (defined by Eqs. 3 and 4, Methods), the waveform of inhibitory firing rates and synaptic gating also closely followed injected EPSCs. As increased input to inhibitory populations leads to increased inhibitory effect and vice versa, resulting feedback inhibition (i.e. IPSC) waveforms were inverted to injected EPSCs. In other words, excitation and inhibition were balanced during each cycle, which is in accordance with published electrophysiology results (Atallah & Scanziani, 2009; Okun & Lampl, 2008). Consequently, IPSCs peaked during the trough of the α-phase and were lowest during the peak of the α-phase. Fitting the models to fMRI activity resulted in a biologically plausible ratio (Atallah & Scanziani, 2009; Xue et al., 2014) of EPSCs to IPSCs, with IPSC amplitudes being about three times larger than EPSC amplitudes (Figure 5b). Because IPSCs have dominated excitatory population inputs, firing rates of excitatory populations showed a similar shape as feedback inhibition currents, i.e., they peaked during the trough of the α-cycle and fell to their minimum during the peak of the α-cycle, reproducing their empirical relationship (Haegens et al., 2011).

In summary, the fast population activity underlying fMRI predictions showed a rhythmic modulation of firing rates on the fast time scale of individual α-cycles in accordance with empirical observations (Haegens et al., 2011). Analyses revealed that periodically alternating states of excitation and inhibition resulted from the ongoing balancing of EPSCs by feedback IPSCs, which explains α-phase related neural firing.

### α-power fluctuations generate fMRI oscillations

Similar to intracranial recordings in monkey (Haegens et al., 2011), we found that increased alpha power of injected EEG source activity was accompanied by decreased firing rates (Figure 6—figure supplement 1). Furthermore, we also observed the empirically observed inverse relationship between α-power and fMRI amplitude (Goldman et al., 2002; Moosmann et al., 2003) in our empirical data in the form of negative correlations between the α-regressor and fMRI activity (Figure 3). Our findings raised the question what physiological mechanism led to this inverse relationship between α-power, firing rate, and respectively fMRI amplitude. We therefore analysed model activity on the longer time scale of α-power fluctuations. To isolate the effects of α-waves from other EEG rhythms, we replaced the injected EEG-source activity in the 15 individual hybrid models with artificial α-activity and simulated all 15 hybrid models using the single parameter set that previously generated the highest average fMRI time series prediction quality (Figure 3—figure supplement 1). Injected activity consisted of a 10 Hz sine wave that contained a single brief high power burst in its centre in order to allow for model activity to stabilize for sufficiently long phases before and after the high power burst. After simulation we computed grand average waveforms of model state variables over all simulated region time series and found that input currents, firing rates, synaptic activity and fMRI activity of excitatory populations decreased in response to the α-burst (Figure 6a). Notably, this behaviour emerged despite the fact that injected activity was cantered at zero, i.e., positive and negative deflections of input currents were balanced. The reason for the observed asymmetric response to increasing input α-power levels originated from inhibitory population dynamics: while positive deflections of α-cycles generated large peaks in ongoing firing rates of inhibitory populations, negative deflections were bounded by 0 Hz. Because of this rectification of high-amplitude negative half-cycles, average per-cycle firing rates of inhibitory populations increased with increasing α-power. As a result, also feedback inhibition had increased for increasing α-power, which in turn led to increased inhibition of excitatory populations, decreased average firing rates, synaptic gating variables and ultimately fMRI amplitudes.

We next analysed the relationship between α-power fluctuations and fMRI oscillations. We generated artificial α-activity consisting of a 10 Hz sine wave that was amplitude modulated by slow oscillations (cycle frequencies between 0.01 and 0.03 Hz) and injected it into the hybrid models of all subjects (Figure 6b). As in the previous example, inhibitory populations filtered negative α-deflections during epochs of increased power. This half-wave rectification led to a modulation of average per-cycle firing rates in proportion to α-power, which introduced a new slow frequency component into the resulting time series. The activity of inhibitory populations can be compared to envelope detection used in radio communication for AM signal demodulation. The new frequency component introduced by half-wave rectification of α-activity modulated feedback inhibition, which in turn modulated excitatory population firing rates. Furthermore, the resulting oscillation of firing rates was propagated to synaptic dynamics where the large time constant of NMDAergic synaptic gating *(τ_NMDA_* = 100 ms vs. ***τ_GABA_*** = 10 ms) led to an attenuation of higher frequencies. The low-pass filtering property of the hemodynamic response additionally attenuated higher frequencies such that in fMRI signals only the slow frequency components remained. To restate: α-power fluctuation introduced an inverted slow modulation of firing rates and synaptic activity; the low-pass filtering properties of synaptic gating and hemodynamic responses attenuated higher frequencies such that only the slow oscillation remained in fMRI signals. To check whether this mechanism is robust to the choice of the frequency of the injected alpha rhythm (10 Hz) we simulated otherwise identical models for artificial alpha waves at 9 Hz and 11 Hz frequencies and found qualitatively identical results: simulated fMRI and moving average firing rate time series of the 9 Hz and the 11 Hz model had correlation coefficients *r* > 0.99 with the respective time series of the 10 Hz model.

In summary, we found that increased α-power led to increased feedback inhibition of excitatory populations introducing a slow modulation of population firing, which can explain the empirically observed anticorrelation between α-power and fMRI.

### Long-range coupling controls fMRI power-law scaling

Empirical fMRI power spectra follow a power-law distribution *P ∝f^β^*, where *P* is power, *f* is frequency and *β* the power-law exponent, which is an indicator of criticality and scale-free dynamics that appears throughout nature (Beggs & Timme, 2012; He, 2011). In accordance with systematic analyses of empirical data (He, 2011), average power spectra of our empirical fMRI data obeyed power-law distributions with exponent *β_emp_* =-0.82 (Figure 7a and Figure 7—figure supplement 1). Time series were tested for scale invariance using rigorous model selection criteria that overcome the limitations of simple straight-line fits to power spectra for estimating scale invariance (see Methods; for illustration purposes straight-line fits are shown in Figure 7a and Figure 7—figure supplement 1).

Our previous results associated resting-state fMRI oscillations with EEG by identifying a neural mechanism that transforms instantaneous EEG source power fluctuations into fMRI oscillations (Figure 6). Surprisingly, however, the power spectrum of EEG source power had a considerably smaller negative exponent than fMRI *(β_α-band_* =-0.53 for α-power and *β_wide-band_* =-0.47 for wide-band power). Comparison of power spectra indicated that power-law fits of simulated fMRI power spectra had a higher negative exponent than power-law fits of source-activity power spectra because the power of slower oscillations increased relative to the power of faster oscillations (Figure 7a and Figure 7—figure supplement 1). That is, model dynamics transformed input activity such that the amplitude of output oscillations increased inversely proportional to their frequency. The effect is visible in Figure 6b, where fMRI, synaptic and firing rate amplitudes of slow oscillations were larger than amplitudes of fast oscillations, despite equally large amplitudes of input α-power oscillations. In comparison, simulation results obtained for deactivated large-scale coupling, but an otherwise identical setup, did not show this frequency-dependent amplification (Figure 7b). Without large-scale coupling the power-law exponent of simulated fMRI *(β_sim_Gzero_* =-0. 54) was close to the exponent of alpha power time series of injected EEG source activity *(β_α-band_* =-0.53).

Comparison of the individual components of population inputs for activated (Figure 6b, column II) vs. deactivated (Figure 7b, column II) long-range coupling show that the only difference between both setups is the shape of long-range input in the former case. When long-range coupling is activated, the emerging long-range input (column II, green trace) shows in-phase coherence with IPSCs (column II, blue trace, moving average: black trace), which leads to constructive interference of both waves and in turn to a stronger modulation of total excitatory population inputs compared to the deactivated case (column III, red trace, moving average: black trace). This constructive interference of long-range input and IPSC oscillations in combination with slowly decaying NMDA synaptic gating activity (Eq. 5, effective time constant of NMDA is **τ_NMDA_**= 100 ms) leads to a situation in which NMDA synaptic gating activity accumulates. The period of time for which this excitatory feedback persists is longer during slower oscillations than during faster oscillations. Consequently, synaptic activity (column VI) has more time to accumulate and is therefore larger during slower oscillations compared to faster oscillations. As a result, the amplitudes of excitatory population fMRI (column VII, red traces) reach higher values during slower oscillations than during faster oscillations for activated large-scale coupling (Figure 6b). Accordingly, the power of slower oscillations, and therefore the slope of the power spectrum, increases in the case of long-range coupling. Note that this effect (i.e., that slower oscillations reach higher amplitudes) can already be observed in firing rates and synaptic gating time series, which excludes an influence of the hemodynamic forward model. In contrast, in the case of deactivated large-scale coupling (Figure 6b) all amplitude peaks are approximately equal, which was the expected result, since the amplitude-peaks of the power modulation of injected α-activity were equally high by construction (column I, orange trace).

We asked how the relative strengths of white-matter excitation and feedback inhibition influence scale-free dynamics. In order to test how E/I balance affects power-law scaling of neural activity, we varied the parameters that control the strength of global white-matter coupling and global feedback inhibition (the latter being controlled by a single parameter for all inhibitory populations), while keeping the strengths of EEG source activity injected into excitatory and inhibitory populations fixated. Screening of individual parameter spaces showed that the power-law exponent of simulated fMRI depended on the balance of large-scale excitation and local inhibition: the 2D distribution of prediction quality (of fMRI time series, functional connectivity and the power-law exponent) showed a characteristic diagonal pattern, which demonstrated the crucial role of E/I balance for the emergence of scale invariance and long-range correlations Figure 7—figure supplement 1 and Figure 7—figure supplement 2).

## Discussion

In this work we describe a biophysically based brain network model that predicts a considerable part of ongoing subject-specific fMRI resting-state time series. Furthermore, we show how this novel modelling approach can be used to infer the neurophysiological mechanisms underlying neuroimaging signals. Instead of mere reproduction of empirical observations, our central aim was to provide an integrative framework that unifies empirical data with theory of the nervous system in order to derive mechanisms of brain function underlying empirical observations across many scales. A key point of consideration is that the brain model was built from networks of generic neural population models that were constrained by empirical data, but not explicitly constructed to address specific reproduced phenomena. This is mirrored by the emergence of processes at considerably faster time scales than the subject-specific fMRI time series that were the target of the model fitting. It is important to point out that the inferred mechanisms constitute candidate hypotheses that require empirical falsification. The model-derived mechanisms make concrete predictions on the waveforms of different input currents, output firing rates, synaptic activities and fMRI signals, which can be empirically tested. Through ongoing integration of biological knowledge, falsification with empirical data and subsequent refinement, hybrid brain network models are intended to represent a comprehensive and increasingly accurate theory about large-scale brain function. The construction of hybrid brain network models and our major results are visualized in **Supplementary Movie 1**.

Hybrid models draw on empirically estimated EEG source activity to constrain input current dynamics. Models, by definition, omit features of the modelled system for the sake of simplicity, generality and efficiency. Adding degrees of freedom renders parameter spaces increasingly intractable and increases the risk of over-fitting. Injection of source activity is a way to systematically probe sufficiently abstract neural systems while maintaining biologically realistic behaviour. Thereby, the approach aims to balance a level of abstraction that is sufficient to provide relevant insights, with being detailed enough to guide subsequent empirical study. It is not the goal of this approach to attain the highest possible fit between different imaging modalities at the cost of biological plausibility, which would be the case for abstract statistical models that do not relate to biological entities and therefore preclude the inference of neurophysiological knowledge. Here, imperfect reproduction of neural activity directly points to deficits in our understanding and conceptualization of large-scale brain structure and function, which to iteratively improve is the goal of this approach.

In line with our results, cellular-level studies indicate that rhythmic GABAergic input from the interneuronal network is associated with E/I balance (Dehghani et al., 2016) and α-related pulsed inhibition (Jensen & Mazaheri, 2010; Lőrincz et al., 2009; Osipova et al., 2008). However, the identification of an exact physiological mechanism that explains how α-rhythms can produce an inhibitory effect remained elusive (Jensen & Mazaheri, 2010; Klimesch, 2012). Mazaheri et al. (Mazaheri & Jensen, 2010) suggest that pulsed inhibition occurs due to an observed amplitude asymmetry of ongoing oscillations, also termed baseline-shift. Our results suggest, in accordance with the model from Mazaheri et al. (Mazaheri & Jensen, 2010), that a symmetrically oscillating driving signal in the α-range leads to asymmetric firing rates and synaptic currents, but we extend this scheme with an explicit explanation of the generation of inhibitory pulses from oscillating input currents.

It is unclear to which degree non-neuronal processes affect the fMRI signal, as different physiological signals such as respiration and cardiac pulse rate were shown to be correlated with resting-state oscillations (Biswal et al., 1996; Power et al., 2016), which raised concerns that RSN oscillations may be unrelated to neuronal information processing, but rather constitute an epiphenomenon (Birn et al., 2006; de Munck et al., 2008; Shmueli et al., 2007; Yuan et al., 2013). The interpretation and handling of these signal modulations is therefore hotly debated and they are often considered as artifactual and removed from fMRI studies (Birn et al., 2006; Chang & Glover, 2009). Importantly, however, low-frequency

BOLD fluctuations are also strongly correlated with electrical neural activity, which was shown by studies that analysed fMRI jointly with EEG (Goldman et al., 2002; Laufs et al., 2003; Moosmann et al., 2003), intracortical recordings (He et al., 2008; Logothetis et al., 2001) or MEG (Brookes et al., 2011; de Pasquale et al., 2010). Similarly, strong temporal correlations and spatially similar correlation maps of EEG α-power, respiration and BOLD (Yuan et al., 2013), as well as of EEG α-power, heart rate variations and BOLD (de Munck et al., 2008) suggest that these fMRI fluctuations are not unrelated to neural activity, but may be of neural origin.

Our results extend the current understanding by showing an explicit mechanism for a neural origin of fMRI RSN oscillations that explains a large part of their variance by a chain of neurophysiological interactions. That is, our simulated activity not only reproduces the negative correlation between α-power fluctuations and BOLD signal, but also reveals a mechanism that transforms ongoing α-power fluctuation into fMRI oscillations. In addition to fMRI time series, the hybrid model also reproduces the spatial topology of fMRI networks, which are not predicted by the α-power regressor. These findings thereby add to accumulating evidence suggesting that RSNs originate from neuronal activity (Brookes et al., 2011; de Pasquale et al., 2010; Goldman et al., 2002; He et al., 2008; Logothetis et al., 2001; Mantini et al., 2007; Moosmann et al., 2003) rather than being a purely hemodynamic phenomenon that is only correlated, but not caused by it (Birn et al., 2006; de Munck et al., 2008; Shmueli et al., 2007). The conclusions from these results have important implications for future fMRI studies, as they implicate that low-frequency fMRI oscillations may be attributed to a neural process that has a considerable state-dependent effect on neural information processing as indicated by the large modulations of neuronal firing and synaptic activity. Methods for physiological noise correction might remove variance from fMRI experiments that is related to neuronal activity and may therefore exclude relevant information for the interpretation of fMRI data.

Parameter space exploration shows that structural coupling is critical for fMRI prediction, as prediction quality decreases for sub-optimal global coupling strengths or when global coupling is deactivated altogether (Figure 7—figure supplement 2). We observed that the prediction quality of resting-state network activation time courses fluctuates over time and is highest during epochs of highest variance of the respective temporal mode. During these time windows resting-state networks contribute the largest variance to whole-brain fMRI, i.e., they are the most active. A possible explanation may be that during states of asynchronous neural activity (i.e. states of low functional network activity) destructive interference of electromagnetic waves decreases the ability of source imaging methods to reconstruct source activity. It is important to note that the observed processes may not be specific to α-oscillations, but may apply also to other frequencies or non-oscillatory signal components. Additional empirical and theoretical studies will be needed to address these limitations more comprehensively.

Despite the ubiquity of scale invariant dynamics, models that generate power-law distributions are often rather generic and detached from the details of the modelled systems (Bak et al., 1987; Marković & Gros, 2014). Furthermore, the precise mechanisms that lead to the emergence of fMRI power spectrum power-law scaling or the relationship between brain network interaction and fMRI power-law scaling are unclear (He, 2011). Our simulation results indicate that fMRI spectra power-law scaling is due to the observed frequency-dependent amplification of oscillatory activity in networks that contain self-reinforcing feedback excitation together with slow decay of activity. Central to theories on the emergence of criticality is the tuning of a control parameter (e.g. connection strengths) that leads the system to a sharp change in one or more order parameters (e.g. firing rates) when the control parameter is moved over a critical point that marks the boundary of a phase transition. *In vivo, in vitro* and *in silico* results show that the dynamical balance between excitation and inhibition was found to be essential to move the system towards or away from criticality, e.g., by pharmacologically altering the excitation-inhibition balance in anesthetized rats (Osorio et al., 2010), acute slices (Beggs & Plenz, 2003) or by changing parameters that control global excitation and inhibition in computational models (Deco et al., 2014). However, the exact role played by excitation-inhibition balance is unclear. In line with these results, we found that power-law scaling varied as a function of the relative levels of global excitation and inhibition, further emphasizing the need for a proportional relationship between these control parameters (Figure 7—figure supplement 2). Extending from that, our simulation results indicate that E/I balance may cause a tuning of the relative strengths of local and long-range inputs to neural populations that supports constructive interference between the different input currents, which in turn amplifies slower oscillations more than faster oscillation. These results address an open question on whether power-laws in neural networks result from power-law behaviour on the cellular level or from a global network-level process (Beggs & Timme, 2012), by giving an explanation for scale-free fMRI power spectra as an emergent property of large-scale brain network interaction that does not require small-scale decentralized processes like the constant active retuning of microscopic parameters as proposed in some theories of self-organized criticality (Bak et al., 1987; Hesse & Gross, 2015). Furthermore, these results explicitly address the effect of input activity, while *in vitro* and *in silico* studies have so far focused on systems without or considerably decreased input (Hesse & Gross, 2015). The observed co-emergence of spatial long-range correlations (i.e. functional connectivity networks) and power-law scaling may point to a unifying explanation within the theory of self-organized criticality, as previously proposed by others (Linkenkaer-Hansen et al., 2001).

A wide range of disorders like autism, schizophrenia, intellectual disabilities, Alzheimer’s disease, multiple sclerosis or epilepsy have been linked to disruption of E/I balance (Marín, 2012) and altered structural and functional network connectivity (Stam, 2014). The presented modelling approach may therefore play a key role for identifying the precise mechanisms underlying the pathophysiology of different disorders and assist in developing novel therapies that restore altered E/I balance or brain connectivity, e.g., by identifying the targets for neural stimulation therapies or by guiding individually customized therapy.

## Methods

### Computational model

The model used in this study is based on the large-scale dynamical mean field model used by Deco and colleagues (Deco et al., 2014; Wong & Wang, 2006). Brain activity is modelled as the network interaction of local population models that represent cortical areas. Cortical regions are modelled by interconnected excitatory and inhibitory neural mass models. In contrast to the original model, excitatory connections were replaced by injected EEG source activity. The dynamic mean field model faithfully approximates the time evolution of average synaptic activities and firing rates of a network of spiking neurons by a system of coupled non-linear differential equations for each node *i*:

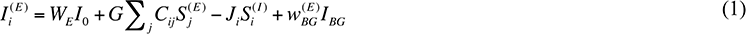

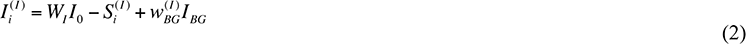

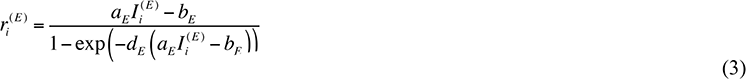

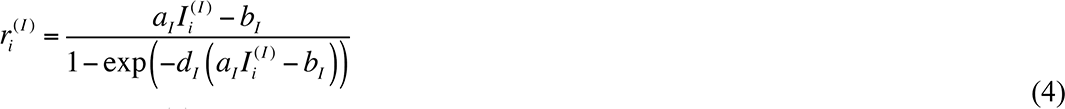

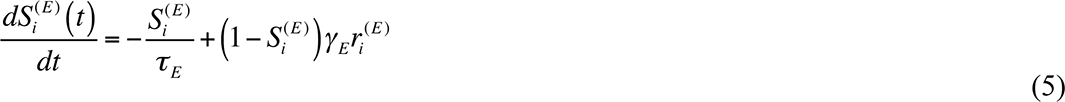

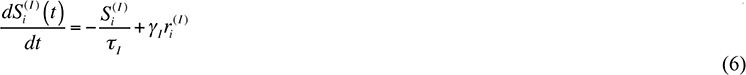

Here, 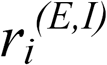 denotes the population firing rate of the excitatory *(E)* and inhibitory *(I)* population of brain area *i*. 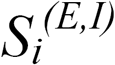 identifies the average excitatory or inhibitory synaptic gating variables of each brain area, while their input currents are given by 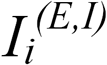. In contrast to the model used by Deco et al.(Deco et al., 2014) that has recurrent and feedforward excitatory coupling, we approximate excitatory postsynaptic currents *IBG* using region-wise aggregated EEG source activity that is added to the sum of input currents 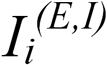. This approach is based on intracortical recordings that suggest that EPSCs are non-random, but strongly correlated with electric fields in their vicinity, while IPSCs are anticorrelated with EPSCs^8-10^. The weight parameters ω_BG_^(E,I)^rescale the z-score normalized EEG source activity independently for excitatory and inhibitory populations. *G* denotes the large-scale coupling strength scaling factor that rescales the structural connectivity matrix *C_ij_* that denotes the strength of interaction for each region pair *i* and *j*. All three scaling parameters are estimated by fitting simulation results to empirical fMRI data by exhaustive search. Initially, parameter space (n-dimensional real space with *n* being the number of optimized parameters) was constrained such that the strength of inhibition was larger than the strength of excitation, satisfying a biological constraint. Furthermore, for each tested parameter set (containing the three scaling parameters mentioned above) the region-wise parameters *J_i_* that describe the strength of the local feedback inhibitory synaptic coupling for each area *i* (expressed in nA) are fitted with the algorithm described below such that the average firing rate of each excitatory population in the model was close to 3.06 Hz (i.e. the cost function for tuning parameters *J_i_* was solely based on average firing rates and not on prediction quality). The overall effective external input I_0_=0.382 nA is scaled by *W_E_* and *W_I_*, for the excitatory and inhibitory pools, respectively. 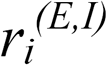 denotes the neuronal input-output functions (f-I curves) of the excitatory and inhibitory pools, respectively. All parameters except those that are tuned during parameter estimation are set as in Deco et al. (Deco et al., 2014). BOLD activity was simulated on the basis of the excitatory synaptic activity *S^(E)^* using the Balloon-Windkessel hemodynamic model(Friston et al., 2003).

### Parameter optimization

For each brain network model, three parameters were varied to maximize the fit between empirical and simulated fMRI: the scaling of excitatory white-matter coupling and the strengths of the inputs injected into excitatory and inhibitory populations. Following *in vivo* observations (Atallah & Scanziani, 2009; Xue et al., 2014), we ensured the EPSC amplitudes at excitatory populations are smaller than IPSC amplitudes by constraining parameters such that the standard deviation of injected source activity was larger for inhibitory populations, i.e., *ω_BG_^(E)^<ω_BG_^(I)^*. Specifically, we scanned 12 different ratios of both parameters *ω_BG_^(I)^* / *ω_BG_^(E)^* with values between 5 and 200. Besides tuning these three global parameters, which were the sole optimization criterion to maximize the fit of simulated activity with empirical fMRI time series, we adjusted local inhibitory coupling strengths in order to obtain biologically plausible firing rates in excitatory populations. For this second form of tuning, termed feedback inhibition control (FIC), average population firing rates were the sole optimization criterion, without any consideration of prediction quality, which was only dependent on the three global parameters. FIC modulates the strengths of inhibitory connections that is required to compensate for excess or lack of excitation resulting from the large variability in white-matter coupling strengths obtained by MRI tractography, which is a prerequisite to obtain plausible ranges of population activity that is relevant for some results (Figure 5 and Figure 6). Prediction quality was measured as the average correlation coefficient between all simulated and empirical region-wise fMRI time series of a complete cortical parcellation over 20.7 minutes length (TR = 1.94s, 640 data points) thereby quantifying the ability of the model to predict the activity of 68 parcellated cortical regions. Accounting for the large-scale nature of fMRI resting-state networks, the chosen parcellation size provides a parsimonious trade-off between model complexity and the desired level of explanation. What this parcellation may lack in spatial detail, it gains in providing a full-brain coverage that can reliably reproduce ubiquitous large-scale features of empirical data, which we further present below. To exclude overfitting and limited generalizability, a five-fold cross-validation scheme was performed on the hybrid model simulation results. Therefore, the data was randomly divided into two subsets: 80 % as training subset and 20 % as testing subset. Prediction quality was estimated using the training set, before trained models were asked to predict the testing set. Resulting prediction quality was compared between training and test data set and between test data set and the data obtained from fitting the full time series. Furthermore, despite the large range of possible parameters, the search converged to a global maximum (Figure 3—figure supplement 1). Therefore, we ensured that when the model has been fit to a subset of empirical data, that it was able to generalize to new or unseen data. In contrast to model selection approaches, where the predictive power of different models and their complexity are compared against each other, we here use only a single type of model.

### Feedback inhibition control

Using standard parameters of the original model, the excitatory populations of isolated nodes have an average firing rate of 3.06 Hz, which conforms to the Poisson-like cortical *in vivo* activity of ~3 Hz (Softky & Koch, 1993; Wilson et al., 1994). For coupled populations, firing rates change in dependence of the employed structural connectivity matrix and injected input. To compensate for a resulting excess or lack of excitation, a local regulation mechanism, called feedback inhibition control (FIC), was used. The approach was previously successfully used to significantly improve FC prediction as well as for increasing the dynamical repertoire of evoked activity and the accuracy of external stimulus encoding (Deco et al., 2014). Despite the mentioned advantages of FIC tuning, it has the disadvantage of increasing the number of open parameters of the model. To prove that prediction quality is not due to FIC, but solely due to the three global parameters and to exclude concerns about over-parameterization or that FIC may be a potentially necessary condition for the emergence of scale-freeness, we devised a control model that did not implement FIC, but used a single global parameter for inhibitory coupling strength. Instead of tuning the 68 individual local coupling weights individually, only a single global value for all inhibitory coupling weights *J_i_* was varied. We compared the effect of FIC on time series prediction quality and found no significant difference in prediction quality to simulations that used only a single value for all local coupling weights *J_i_* per subject (one-tailed Wilcoxon rank sum test, *p* = 0.36, *z* =-0.37, Cohen’s *d* =-0.038). In contrast to simulations that are driven by noise (Deco et al., 2014), FIC parameters for injected input must be estimated for the entire simulated time series, since the non-stationarity of stimulation time series leads to considerable fluctuations of firing rates. Therefore, we developed a local greedy search algorithm for fast FIC parameter estimation based on the algorithm in Deco et al. (Deco et al., 2014). To exert FIC, local inhibitory synaptic strength is iteratively adjusted until all excitatory populations attained a firing rate close to the desired mean firing rates for the entire ~20 minutes of activity. During each iteration, the algorithm performs a simulation of the entire time series. Then, it computes the mean firing activity over the entire time series for each excitatory population and adapts *J_i_* values accordingly, i.e., it increases local *J_i_* values if the average firing rate over all excitatory populations during the k-th iteration 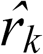 is larger than 3.6Hz and vice versa. In order to reduce the number of iterations the value by which *J_i_* is changed is, in contrast to the algorithm by Deco et al. (Deco et al., 2014), dynamically adapted in dependence of the firing rate obtained during the current iteration

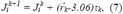

where 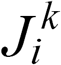 denotes the value of feedback inhibition strength of node *i* and *τ_k_* denotes the adaptive tuning factor during the k-th iteration. In the first iteration, all *J_i_* values are initialized with 1 and *τ_k_* is initialized with 0.005. The adaptive tuning factor is dynamically changed during each iteration based on the result of the previous iteration:

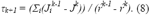

For the case that the result did not improve during the current iteration, i.e.,

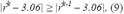

the adaptive tuning factor is decreased by multiplying it with 0.5 and the algorithm continues with the next iteration. After 12 iterations, all *J_i_* values are set to the values they had during the iteration *k* where 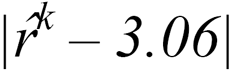 was minimal.

### MRI preprocessing

Structural and functional connectomes from 15 healthy human subjects (age range: 18-31 years, 8 female) were extracted from full data sets (diffusion-weighted

MRI, T1-weighted MRI, EEG-fMRI) using a local installation of a pipeline for automatic processing of functional and diffusion-weighted MRI data (Schirner et al., 2015). From a local database of 49 subjects (age range 18-80 years, 30 female) that was acquired for a previous study (Schirner et al., 2015) we selected the 15 youngest subjects that fulfilled highest EEG quality standards after applying MR artifact correction routines. EEG quality was assessed by standards that were defined prior to the experimental design and that are routinely used in the field (Becker et al., 2011; Freyer et al., 2009; Ritter et al., 2010; Ritter et al., 2007): occurrence of spikes in frequencies >20 Hz in power spectral densities, excessive head motion and cardio-ballistic artifacts. Research was performed in compliance with the Code of Ethics of the World Medical Association (Declaration of Helsinki). Written informed consent was provided by all subjects with an understanding of the study prior to data collection, and was approved by the local ethics committee in accordance with the institutional guidelines at Charité Hospital Berlin. Subjects with a self-reported history of neurological, cognitive, or psychiatric conditions were excluded from the experiment. Structural (T1-weighted high-resolution three-dimensional MP-RAGE sequence; TR = 1,900 ms, TE = 2.52 ms, TI = 900 ms, flip angle = 9°, field of view (FOV) = 256 mm x 256 mm x 192 mm, 256 x 256 x 192 Matrix, 1.0 mm isotropic voxel resolution), diffusion-weighted (T2-weighted sequence; TR = 7,500 ms, TE = 86 ms, FOV = 192 mm x 192 mm, 96 x 96 Matrix, 61 slices, 2.3 mm isotropic voxel resolution, 64 diffusion directions), and fMRI data (two-dimensional T2-weighted gradient echo planar imaging blood oxygen level-dependent contrast sequence; TR = 1,940 ms, TE = 30 ms, flip angle = 78°, FOV = 192 mm x 192 mm, 3 mm x 3 mm voxel resolution, 3 mm slice thickness, 64 x 64 matrix, 33 slices, 0.51 ms echo spacing, 668 TRs, 7 initial images were acquired and discarded to allow magnetization to reach equilibrium; eyes-closed resting-state) were acquired on a 12-channel Siemens 3 Tesla Trio MRI scanner at the Berlin Center for Advanced Neuroimaging, Berlin, Germany. Extracted structural connectivity matrices intend to give an aggregated representation of the strengths and time-delays of interaction between regions as mediated by white matter fiber tracts. As for the original model by Deco et al. (Deco et al., 2014), conduction delays were neglected in this study as it was shown previously that they can be neglected for region-level simulations and activity that is slower than oscillations in the gamma range (Proix et al., 2016). Strength matrices *C_ij_* were divided by their respective maximum value for normalization. In short, the pipeline proceeds as follows: for each subject a three-dimensional high-resolution T1-weighted image image was used to divide cortical gray matter into 68 regions according to the Desikan-Killiany atlas using FreeSurfer’s (Fischl, 2012) automatic anatomical segmentation and registered to diffusion data. The gyral-based brain parcellation is generated by an automated probabilistic labeling algorithm that has been shown to achieve a high level of anatomical accuracy for identification of regions while accounting for a wide range of inter-subject anatomical variability (Desikan et al., 2006). The atlas was successfully used in previous modelling studies and provided highly significant structure-function relationships (Honey et al., 2009; Ritter et al., 2013; Schirner et al., 2015). Details on diffusion-weighted and fMRI preprocessing can be found in Schirner et al. (Schirner et al., 2015) Briefly, probabilistic white matter tractography and track aggregation between each region-pair was performed as implemented in the automatic pipeline and the implemented distinct connection metric extracted. This metric weights the raw track count between two regions according to the gray-matter/white-matter interface areas of both regions used to connect these regions in distinction to other metrics that use the unweighted raw track count, which was shown to be biased by subject-specific anatomical features (see Schirner et al. (Schirner et al., 2015) for a discussion). After preprocessing, the cortical parcellation mask was registered to fMRI resting-state data of subjects and average fMRI signals for each region were extracted. The first five images of each scanning run were discarded to allow the MRI signal to reach steady state. To identify RSN activity a spatial Group ICA decomposition was performed for the fMRI data of all subjects using FSL MELODIC (Beckmann & Smith, 2004) (MELODIC v4.0; FMRIB Oxford University, UK) with the following parameters: high pass filter cut off: 100 s, MCFLIRT motion correction, BET brain extraction, spatial smoothing 5 mm FWHM, normalization to MNI152, temporal concatenation, dimensionality restriction to 30 output components. ICs that correspond to RSNs were automatically identified by spatial correlation with the nine out of the ten well-matched pairs of networks of the 29,671-subject BrainMap activation database as described in Smith et al. (Smith et al., 2009) (excluding the cerebellum network). All image processing was performed in the native subject space of the different modalities and the brain atlas was transformed from T1-space of the subject into the respective spaces of the different modalities.

### EEG preprocessing

Details of EEG preprocessing are described in supplementary material of Schirner et al. (Schirner et al., 2015). First, to account for slow drifts in EEG channels all channels were high-pass filtered at 1.0 Hz (standard FIR filter). Imaging Acquisition Artefact (IAA) correction was performed using Analyser 2.0 (v2.0.2.5859, Brain Products, Gilching, Germany). The onset of each MRI scan interval was detected using a gradient trigger level of 300 μV/ms. Incorrectly detected markers, e.g. due to shimming events or heavy movement, were manually rejected. To assure the correct detection of the resulting scan start markers each inter-scan interval was controlled for its precise length of 1940 ms (TR). For each channel a template of the IAA was computed using a sliding average approach (window length: 11 intervals) and subsequently subtracted from each scan interval. For further processing, the data was down sampled to 200 Hz, imported to EEGLAB and low-pass filtered at 60 Hz. ECG traces were used to detect and mark each instance of the QRS complex in order to identify ballistocardiogram (BCG) artifacts. The reasonable position and spacing of those ECG markers was controlled by visual inspection and corrected if necessary. To correct for BCG and artifacts induced by muscle activity, especially movement of the eyes, a temporal ICA was computed using the extended Infomax algorithm as implemented in EEGLAB. To identify independent components (ICs) that contain BCG artifacts the topography plot, activation time series, power spectra and heartbeat triggered average potentials of the resulting ICs were used as indication. Based on established characteristics, all components representing the BCG were identified and rejected, i.e., the components were excluded from back-projection. The remaining artificial, non-BCG components, accounting for primarily movement events especially eye movement, were identified by their localization, activation, power spectral properties and ERPs. Detailed descriptions of EEG and fMRI preprocessing have been published elsewhere (Becker et al., 2011; Freyer et al., 2009; Ritter et al., 2010; Ritter et al., 2007).

### Biologically based model input

EEG source imaging was performed with the freely available MATLAB toolbox Brainstorm using default settings and standard procedure for resting-state EEG data as described in the software documentation (Tadel et al., 2011). Source space models were based on the individual cortical mesh triangulations as extracted by FreeSurfer from each subject’s T1-weighted MRI data and downsampled by Brainstorm. From the same MRI data, head surface triangulations were computed by Brainstorm. Standard positions of the used EEG caps (Easy-cap; 64 channels, MR compatible) were aligned by the fiducial points used in Brainstorm and projected onto the nearest point of the head surface. Forward models are based on Boundary Element Method head models computed using the open-source software OpenMEEG and 15002 perpendicular dipole generator models located at the vertices of the cortical surface triangulation. The sLORETA inverse solution was used to estimate the distributed neuronal current density underlying the measured sensor data since it has zero localization error (Pascual-Marqui, 2002). EEG data was low-pass filtered at 30 Hz and imported into Brainstorm. There, the epochs before the first and after the last fMRI scan were discarded and the EEG signal was time-locked to fMRI scan start markers. Using brainstorm routines, EEG data was projected onto the cortical surface using the obtained inversion kernel and averaged according to the Desikan-Killiany parcellation that was also used for the extraction of structural and functional connectomes and region-averaged fMRI signals. The resulting 68 region-wise source time series were imported to MATLAB, z-score normalized and upsampled to 1000 Hz using spline interpolation as implemented by the Octave function *interp1.* To enable efficient simulations, the sampling rate of the injected activity was ten times lower than model sampling rate. Hence, during simulation identical values have been injected during each sequence of ten integration steps.

### Simulation and analysis

Simulations were performed with a highly optimized C implementation of the previously described model on the JURECA supercomputer at the Juelich Supercomputing Center. Simulation and analyses code and used data is open source and available from online repositories (Schirner et al. (2017), see “Data and code availability”). An exhaustive brute-force parameter space scan using 3888 combinations of the parameters *G* and *ω_bg_^(E,I)^* was performed for each subject. Each of these combinations was computed 12 times to iteratively tune *J_i_* values. As control setup, further simulations were performed with random permutations of the input time series. Therefore, each source activity time series was randomly permutated (individually for each region and subject) using the Octave function *randperm()* and injected into simulations using all parameter combinations that were previously used. As an additional control situation the original dynamic mean field model as described in Deco et al. (Deco et al., 2014) was simulated for the 15 SCs. Here, the parameters *G* and *J_NMDA_* were varied and FIC tuning was performed using the same algorithm as used for the source activity injection model. The simulation and FIC optimization process was identical for all three models. The length of the simulated time series for each subject was 21.6 minutes. Simulations were performed at a model sampling rate of 10,000 Hz. BOLD time series were computed for every 10^th^ time step of excitatory synaptic gating activity using the Balloon-Windkessel Model (Friston et al., 2003). From the resulting time series every 1940^th^ step was stored in order to obtain a sampling rate of simulated fMRI that conforms to the empirical fMRI TR of 1.94 s. The first 11 scans (21.34 s) of activity were discarded to allow model activity and simulated fMRI signal to stabilize. For each subject and modelling approach the simulation result that yielded the highest average correlation between all 68 empirical and simulated regions time series for all tested parameters was used for all analyses. To ensure region-specificity of simulation results only corresponding simulated and empirical region time series were correlated in the case of raw fMRI, respectively, for resting-state networks only simulated regions that overlap with the spatial activation pattern of the respective network were used for estimating prediction quality. Specifically, for RSN analysis, only those regions were compared with the temporal modes of RSNs that had a spatial overlap of at least 40 % of all voxels belonging to the respective region. To assess time-varying prediction quality, a correlation analysis was performed in which a window with a length of 100 scans (194 s) was slid over the 68 pairs of empirical and simulated time series and the average correlation over all 68 regions was computed for each window. For the estimation of signal correlation, the computation of entries of FC matrices and as a measure of similarity of FC matrices Pearson’s linear correlation coefficient was used. FC matrices were compared by stacking all elements below the main diagonal into vectors and computing the correlation coefficient of these vectors. Short-term FC prediction quality was estimated by computing the mean correlation obtained for all window-wise correlations of a sliding window analysis of empirical and simulated time series (window-size: 100 scans = 194 s).

To ensure scale-freeness of empirical and simulated signals, region time series were tested using rigorous model selection criteria; on average 79 % of all 1020 region-wise time series (15 subjects x 68 regions) for the seven analysed signal types (empirical fMRI, simulated fMRI, simulated fMRI without global coupling, simulated fMRI without FIC, simulated fMRI without FIC and without global coupling, α-power, α-regressor) tested as scale-free; for every signal type every subject had at least five regions to test as scale-free. PSDs were computed using the Welch method as implemented in Octave, normalized by their total power and averaged. Resulting average power spectra were fitted with a power-law function *f(x) = αx^β^* using least-squares estimation in the frequency range 0.01 Hz and 0.17 Hz which is identical to the range for which the test for scale invariance was performed. Frequencies below were excluded in order to reduce the impact of low-frequency signal confounds and scanner drift, frequencies above that limit were excluded to avoid aliasing artefacts in higher frequency ranges (TR = 1.94 s, hence Nyquist frequency is around 0.25 Hz). In order to compare the scale invariance of our empirical fMRI data with results from previous publications (He, 2011), we also computed power spectra in a range that only included frequencies <0.1 Hz. In order to adequately quantify scale invariance we applied rigorous model selection to every time series to identify power-law scaling and excluded all time series from analyses that were described better by a model other than a power-law. Nevertheless, we compared the obtained results from this strict regime with results obtained when all time series were included and found them to be qualitatively identical. To test for the existence of scale invariance we used a method that combines a modified version of the well-established detrended fluctuation analysis (DFA) with Bayesian model comparison (Ton & Daffertshofer, 2016). DFA is, in contrast to PSD analyses, robust to both stationary and nonstationary data in the presence of confounding (weakly non-linear) trends. Rather than averaging the mean squared fluctuations over consecutive intervals as in conventional DFA, this method uses the values per interval to approximate the distribution of mean squared fluctuations with kernel density estimation. This allows for estimating the corresponding log-likelihood as a function of interval size without presuming the fluctuations to be normally distributed, as in the case of conventional DFA and therefore gives a non-parametric estimate of the log-likelihood for fitted models. Furthermore, conventional DFA does not provide any means to determine whether a power law is present or not. It is important to note, that a simple linear fit of the detrended fluctuation curve without proper comparison of the obtained goodness of fit with that of other models would entirely ignore alternative representations of the data different than a power law. For quantification of the goodness of fit with simple regression its corresponding coefficient of determination, *R^2^*, is ill-suited as it measures only the strength of a linear relationship and is inadequate for nonlinear regression (Ton & Daffertshofer, 2016). Here, we assess power-law scaling in the context of DFA, i.e. the optimality of a straight line fit of fluctuation magnitude against interval size in a log-log representation, with non-parametric model selection using the Bayesian information criterion in order to compare the linear model against alternative models. It is important to note that with this method the assessment of power-law scaling is based on maximum likelihood estimation, which overcomes the limitations of a minimal least-squares estimate obtained from linear regression in the conventional DFA approach. Details of the used method can be found elsewhere (Ton & Daffertshofer, 2016). Briefly, the method first estimates the cumulative sum of each time series. Next, signals are divided into non-overlapping intervals of increasing length, for a range of window sizes (48 time windows in steps from 3 to 50 data points). Then, for each interval the linear trend is removed and the root mean squared (RMS) fluctuation for each detrended interval is computed. Interval size is plotted against the RMS magnitude of fluctuations in a log-log representation and model-fits with 11 different models are computed using maximum likelihood estimation. A straight line on the log-log graph indicates scale invariance expressed as *F(n) ∝ n^α^*, with *n* the interval size, *F(n)* detrended fluctuation and a, the scaling exponent, which represents the slope of the straight line fit (α ≅ 0.5, indicates uncorrelated white noise, while in the case of *α* ≥ 0.5 the auto-correlation function decays slower than the auto-correlation function of Brownian motion, indicating long-term ‘memory’). Lastly, likelihoods are used to compute Bayesian information criteria (BIC) for each model, which are used to select among models. BIC take into account model accuracy (as quantified by maximum likelihood) and model complexity, which scores the number of free parameters used in the different models. Optimality in the context of BIC therefore yields a maximally accurate while minimally complex explanation for data, i.e., the optimal compromise between goodness-of-fit and parsimony. For the different signals the majority of time series were tested as being scale free: 83 % for empirical fMRI, 69 % for simulated fMRI, 71 % for simulated fMRI with deactivated FIC, 83 % for simulated fMRI with deactivated global coupling, 86 % for simulated fMRI with deactivated global coupling and FIC, 90 % for α-power and 70 % for the α-regressor.

To compute grand average waveforms, state-variables were averaged over all 15 subjects and 68 regions (N = 1020 region time series) time-locked to the zero crossing of the α-amplitude, which was obtained by band-pass filtering source activity time series between 8 and 12 Hz; to obtain sharp average waveforms, all α-cycle epochs with a cycle length between 95 and 105 ms were used (N = 4,137,994 α-cycle). For computing ongoing α-power time courses, instantaneous power time series were computed by taking the absolute value of the analytical signal (obtained by the Hilbert transform) of band-pass filtered source activity in the 8-10 Hz frequency range; the first and last ~50 s were discarded to control for edge effects. To compute the α-regressor, power time series were convolved with the canonical hemodynamic response function, downsampled to fMRI sampling rate and shifted relative to fMRI time series to account for the lag of hemodynamic response. The highest negative average correlation over all 68 region-pairs obtained within a range of +/-3 scans shift was used for comparison with simulation results.

### Statistical analyses

All statistical analyses were performed using MATLAB (The MathWorks, Inc., Natick, Massachusetts, United States). Data are represented as box-and-whisker plots. As normality was not achieved for the majority of data sets (assessed by Lilliefors test at significance level of 0.05), differences between groups were compared by non-parametric statistical tests, using either two-tailed Wilcoxon rank sum test or, in case of directional prediction, one-tailed Wilcoxon rank sum test; a value *p* <0.05 was considered significant.

### Data and code availability

Brain network models are implemented in the open source neuroinformatics platform The Virtual Brain that can be downloaded from thevirtualbrain.org. Code and data that support the findings of this study can be obtained from https://github.com/BrainModes/The-Hybrid-Virtual-Brain and https://github.com/BrainModes/The-Hybrid-Virtual-Brain and https://osf.io/mndt8/ (DOI 10.17605/OSF.IO/MNDT8).

## Author contributions

M.S. and P.R. conceived the project. M.S. conceived and implemented the model based on models and code by D.G., V.J. and A.R.M. M.S. assembled input data from previously collected empirical data (Schirner et al., 2015). P.R. supervised the project. M.S. and P.R. analysed and interpreted the results. M.S. wrote the paper with contributions from P.R., A.R.M., V.J. and G.D. All authors discussed the results and commented on the manuscript at all stages.

## Acknowledgments

The authors gratefully acknowledge the computing time granted by the John von Neumann Institute for Computing (NIC) provided on the supercomputer JURECA (Krause & Thörnig, 2016) at Jülich Supercomputing Centre (www.fz-juelich.de, Grant NIC#8344 & NIC#10276 to P.R.). The authors acknowledge the support of the James S. McDonnell Foundation (Brain Network Recovery Group JSMF22002082) to A.R.M., V.J., G.D., and P.R., the German Ministry of Education and Research (US-German Collaboration in Computational Neuroscience, Bernstein Focus State Dependencies of Learning 01GQ0971-5, the Max-Planck Society) and funding from the European Union Horizon2020 (ERC Consolidator grant BrainModes 683049) to P.R. The authors thank Olaf Sporns, Jochen Braun and Andreas Daffertshofer for their helpful comments on the manuscript. The authors declare no competing financial interests.

## Supplementary Movie 1

Reverse-engineering neural information processing. The video shows how computational brain network models are constructed from individual neuroimaging data, how these models can be used to simulate different types of neural activity of individual subjects on multiple temporal scales and how model activity can be used to derive mechanisms of brain function.

**Figure 3—figure supplement 1.**
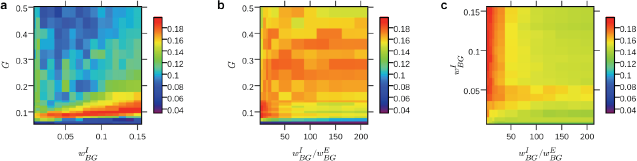
Parameter space exploration results. (**a-c**), 2d parameter space heat maps show average time series correlation over all 68 regions obtained from the hybrid model for different combinations of the three varied parameters scaling of global coupling strength G, scaling of EEG source activity injected into excitatory population ω_BG_^I^, and into inhibitory populations ω_BG_^E^ (the latter depicted as ratio ω_BG_^I^ / ω_BG_^E^); results were averaged over all subjects. Parameter values that yielded the highest average correlation were used for simulations with artificial alpha input (marked with an asterisk). We confirmed identifiability of the model by showing that parameter space search converges towards a single optimal solution yielding best predictions.

**Figure 6—-figure supplement 1.**
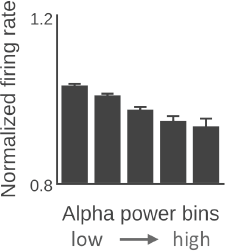
α-power predicts firing rate. In analogy to electrophysiological results shown in Figure 4b in Haegens et al. (Haegens et al., 2011), all time series for each subject and brain region were divided into five equal-sized bins on the basis of α-power level and average firing rate (normalized with average firing rate per brain region) was computed per bin. Firing rate decreased with increasing α-power (median firing rate in all higher-power bins is significantly smaller than in lower-power bins, p <0.001, one-tailed Wilcoxon rank sum test).

**Figure 7—figure supplement 1.**
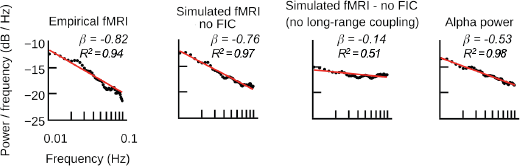
Power spectral densities for simulations with the simplified hybrid model. Power spectra for empirical and simulated fMRI, ongoing alpha power time series of EEG source activity input and the alpha regressor averaged over all subjects and regions (straight-line fits are for illustration purposes only; scale-invariance was determined using rigorous model selection criteria and dynamic fluctuation analysis). For comparison with simulation results shown in Figure 7a, the models used here implemented no feedback inhibition control (FIC), i.e., a single value for all local inhibitory connection parameters J_i_ was used. Empirical and simulated fMRI spectra have a large power-law exponent, i.e. a steeper slope, compared to ongoing alpha power or wide-band power (β_α-band_ =-0.53, β_wide-band_ =-0.47). To analyse the effect of large-scale network interaction, simulated fMRI was computed with and without long-range coupling. To exclude that power-law spectra emerge despite absent large-scale coupling, local inhibition was tuned such that the model produced highest fMRI time series predictions. Without long-range coupling, simulated fMRI showed a similar exponent as injected source activity. When large-scale coupling was activated and global excitation was properly balanced with local inhibition, the exponent was closer to the exponent of empirical fMRI (Figure 7—figure supplement 2).

**Figure 7—figure supplement 2.**
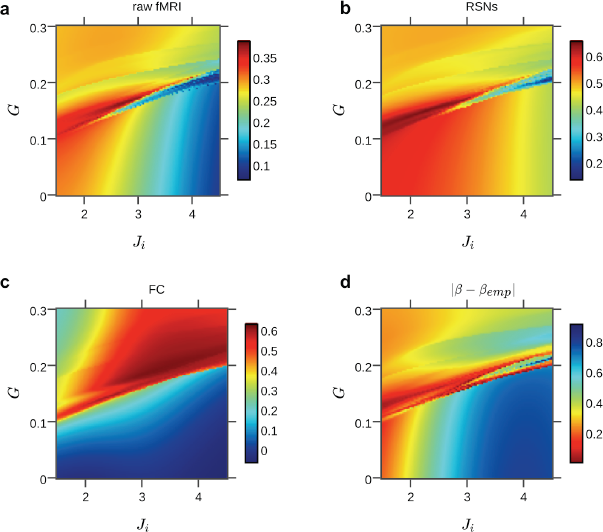
Fine-grained parameter space exploration of the simplified hybrid model. Here, global coupling and a single value for all local inhibitory connection parameters J_i_ was tuned for an exemplary subject. Colors indicate the average correlations between simulated and empirical data (a-c) and the absolute difference between exponents of power-law fits with empirical and simulated power spectral densities (d, colormap was flipped to indicate highest fits in red to be consistently with the other subplots), averaged over all regions. The distributions of the different metrics suggest a link between prediction quality (of raw fMRI, RSNs and FC and the power-law exponent β) and the relative strengths of long-range coupling G and local inhibition J_i_. A diagonal pattern in heat maps indicates that prediction quality and power-law exponents depend on the balancing of large-scale excitation with local inhibition. The plots illustrate that when large-scale coupling was absent or inadequately balanced, models did not produce scale-free behavior and prediction quality decreased.

